# Reshaped functional connectivity gradients in acute ischemic stroke

**DOI:** 10.1101/2024.04.20.590191

**Authors:** Cemal Koba, Joan Falcó-Roget, Alessandro Crimi

## Abstract

Ischemic brain stroke disrupts blood flow, leading to functional and structural changes associated with behavioral deficits. Importantly, despite this disruption occurring in localized regions, the resulting changes in the functional organization are both high-dimensional and widespread across the human cortex. However, the mechanisms with which these global patterns emerge and the subsequent behavioral deficits they entail, remain largely unexplored. Functional connectivity gradients provide consistent, reproducible, and robust low-dimensional representations of brain function that can be explored to reduce brain heterogeneity to a handful of axes along which brain function is organized. Here, we investigated how stroke disrupts this canonical gradient space by aligning each patient to a control-averaged gradient embedding and computing the distances to the “correct” positions to quantify functional deviations and their contribution to behavioral deficits. Importantly, we explicitly corrected these gradients for stroke-induced hemodynamic lags to further study their contribution. We found that lag correction enhanced the functional connectivity gradients most prominently in the second gradient, on which visual and somatomotor function is concentrated. Additionally, we identified significant functional deviations primarily within somatomotor, visual, and ventral attention networks, correlating with behavioral impairments. We studied the hemispheric asymmetries of these deviations finding that intact hemispheres preserve comparable patterns of asymmetry while damaged ones presented important changes. Lastly, right-sided lesions displayed more localized functional deviations than their contralateral lesions. Overall, we provide evidence that 1) correcting for hemodynamic lags improves gradient accuracy, as indicated by increased percentages of explained variance, and 2) behavioral impairments and hemispheric asymmetries result from a repositioning of region-based connectivity profiles in a low-dimensional, interpretable space. This suggests that large-scale brain function alterations manifest in slight, predictable movements largely confined to the visual-somatomotor axis.

## Introduction

Ischemic stroke occurs when a vessel supplying blood to the brain is obstructed, ultimately leading to blood shortage (Roach et al., 2010), causing abnormal neural dynamics even if the blockage is short in time. Importantly, if the impairment lasts long enough, neural death is unavoidable. The infarcted region directly suffering from lack of blood will likely be surrounded by swelling (J. M. Hong et al., 2021), thus extending the structural damage to areas near the onset region. Moreover, large population studies have shown that these infarcted regions are not randomly distributed across the brain, but rather subcortical areas are more likely to be affected (Thiebaut de Schotten et al., 2020; Weaver et al., 2019). Even though a significant heterogeneity concerning the exact location of the ischemia exists, the basal-ganglia networks, the cerebellum, and the cerebrospinal tracts are commonly affected brain areas. Consequently, given the emerging role of such structures in the coordination of distributed and localized neurological functions (Bostan et al., 2013; Quartarone et al., 2020), the diverse findings from functional magnetic resonance imaging (fMRI) studies become more comprehensible.

FMRI crucially relies on the association between the metabolic consumption that neural activity incurs. However, if blood flow is impaired, there is no guarantee that the blood oxygen level-dependent (BOLD) signal reflects artifacts stemming from the stroke or the underlying activity of neural tissue. Crucially, the signal within lesioned areas has been shown to be temporally delayed (Siegel, Snyder, et al., 2016), showing peaks at later time values (Altamura et al., 2009; Bonakdarpour et al., 2015). Furthermore, stroke patients also displayed power spectra that were shifted towards lower frequencies, thus having slower oscillations (Siegel, Snyder, et al., 2016). This empirical delay can be exploited to create perfusion maps solely from the fMRI series (Braban et al., 2023; Khalil et al., 2020; Lv et al., 2013; Tong et al., 2017) by applying temporal shifts so that the cross-correlation between the voxel-wise and a reference signals (e.g., global signal, venous sinuses) is maximum. This *time-lag correction* highlights areas with delayed blood flow, improves the data-quality (Erdoğan et al., 2016), and enhances the functional connectivity in control groups (Tong et al., 2017) while maintaining the relationships with behavioral dysfunctions in stroke studies (Siegel, Snyder, et al., 2016). Importantly, the altered functional topographies of patients are thought to arise from impaired neural function rather than exogenous blood shortage. Nonetheless, time-lag corrections are not ubiquitously practiced (Bayrak et al., 2019), thus adding another layer of heterogeneity to the analyses.

Amongst the wide range of observed behavioral deficits in stroke populations, motor, linguistic, and attention-related are the most common ones (He et al., 2007). For example, during arm-moving tasks, the contralesional hemisphere shows a hyper-activity that, in turn, reduces to baseline levels with the recovery-time (Rehme et al., 2011). Moreover, the degree of this imbalance between ipsi and contralesional hemispheres after post-stroke is descriptive of motor recovery (Dodd et al., 2017). Resting-state (RS) studies broadly converge to a decrease in inter-hemispheric connectivity (i.e., lower Pearson correlations between fMRI series), an increase in connectivity within the contralesional hemisphere (New et al., 2015; Yourganov et al., 2021), and an abnormal functional co-activation between intact and damaged hemispheres (Carter et al., 2010). In general, altered homotopic connectivity predicts lesion size and severity (Tang et al., 2016; Urbin et al., 2014), while specific behavioral deficits are related to damaged connectivity in the corresponding RS network (Carter et al., 2010; Corbetta & Shulman, 2002). But functional data is high-dimensional therefore compromising inferences. Network studies reduce the size of the system by introducing an additional layer of abstraction, where macroscopic regions are used instead of voxels (Schaefer et al., 2018). Although the hub structure of such networks is significantly damaged in stroke patients (Gratton et al., 2012; Zhu et al., 2017), the resulting clinical correlates remain unclear. In addition, intra-class correlation scores are rather low for voxel- and region-wise studies (Noble et al., 2021). Considering these drawbacks for clinical applicability, explicit dimensionality reduction techniques may help to understand the consequences of stroke from a more general point of view. In this regard, functional connectivity gradients (Margulies et al., 2016) display the macroscale functional organization instead of discrete and specific connectivity characteristics between regions or network-based statistics.

Gradients of brain function project the complex functional profiles along three orthogonal axes encapsulating most of their variance (Vos de Wael et al., 2020). Importantly, they have been proven to be highly replicable (Bethlehem et al., 2020; Hardikar et al., 2022; Margulies et al., 2016; Mckeown et al., 2020). In healthy cohorts, the major axis splits the cortex into unimodal and multimodal areas, the second axis separates visual and somatomotor functions, and the third gradient distinguishes active-passive attention modalities (Glasser et al., 2016). But, in clinical groups, slight deformations of these axes were found to be descriptive of specific syndromes, such as autism (S.-J. Hong et al., 2019) and depression (Xia et al., 2022). In stroke, only one study considered this framework to understand the macroscale organization of the brain (dys-)function (Bayrak et al., 2019). Gradient coordinates of voxels within the lesioned sites from the healthy cohort were compared to intact voxels from patients. This yielded a functional distance map that was proportional to the stability of the functional profile over recovery time. Simplifying, voxels that were far from the lesion, in terms of functional similarity, remained largely unaltered through time. However, the data was not corrected for time-lag aberrancies, and masking of the infarcted areas was based on human annotations thus considering the lesion only in an indirect way (i.e., the lesioned signal was discarded). Moreover, these conclusions solely applied to the 1st and 3rd gradients, leaving the visual and somatomotor axis unstudied.

In this 3-dimensional space, each voxel or region of interest (ROI) is uniquely determined by 3 spatial coordinates which, after a correct alignment preserving the original distances between points (Vos de Wael et al., 2020), can be taken as the canonical position in health. We hypothesize that 1) lag-correction repositions each region and increases the alignment with respect to the canonical locations, especially in patients suffering from ischemic stroke, and 2) the position in this space is indicative of functional and behavioral impairments. Although, several metrics have been derived for this purpose (Del Río et al., 2022; Valk et al., 2023), we opt for an interpretable distance function between any two points in space (i.e., euclidean distance between healthy controls and stroke subjects), which we aim to be predictive of behavioral and clinical correlates. We also describe several other known phenomena related to hemispheric differences that affect the functional connectivity gradients.

## Methods

### Dataset details

The dataset was collected by the School of Medicine of the Washington University in St. Louis and complete procedures can be found in Corbetta et al., 2015. They collected MRI data and behavioral examinations of stroke patients and healthy controls. Stroke patients were registered according to the following criteria:

- The subject must be older than 18 years of age,
- The subject must have no history of stroke until the current one,
- The stroke must have up to two lacunes, be clinically silent, be less than 15 mm in size on CT scan,
- Clinical evidence for motor, language, attention, visual, or memory deficits based on neurological examination,
- The onset of stroke must be shorter than 2 weeks during the time of the scan,
- The subject must be awake, alert, and capable of participating in research.

The imaging data consists of structural and functional MRI at three different time points for stroke patients (1-2 weeks, 3 months, and 12 months after stroke), and 2 time points for control patients (initial scan and 3 months later). Structural scans include T1-weighted, T2-weighted, and diffusion tensor images. Functional images include a resting state paradigm. Scanning was performed with a Siemens 3T Tim-Trio scanner at the School of Medicine of the Washington University in St. Louis including structural, functional, and diffusion tensor scans. Structural scans consisted of (1) a sagittal MP-RAGE T1-weighted image (TR=1950 msec, TE=2.26 msec, flip angle=9 deg, voxel size=1.0 x 1.0 x 1.0 mm, slice thickness = 1.00 mm); (2) a transverse turbo spin-echo T2-weighted image (TR=2500 msec, TE=435 msec, voxel-size=1.0 x 1.0 x 1.0 mm, slice thickness = 1.00 mm); and (3) a sagittal FLAIR (fluid-attenuated inversion recovery) (TR=7500 msec, TE=326 msec, voxel-size=1.5 x 1.5 x 1.5 mm, Slice thickness = 1.50 mm). Functional images consisted of 1-7 resting state runs for each session (TR=2000 msec, TE=27 msec, voxel-size= 4 x 4 x 4 mm, slice thickness = 4 mm, total duration = 256 seconds for each run).

### Preprocessing of the imaging data

The dataset was received in DICOM format. Conversion from DICOM to NIFTI format was achieved by an in-house script that operates on Python (Falcó-Roget, 2023). The files were then organized according to the BIDS specifications. In this step, sessions with duplicate scans (back-to-back scanning of any modality) were manually checked by two independent researchers (C.K., J.F.-R.). Scans with lower quality were discarded from future analyses after visual inspection. Multiple scanning happened only when the first acquisition had low quality. Runs that had no T1 or RS scan were removed. The dataset consisted of hemorrhagic and ischemic stroke cases. Only the ischemic stroke cases of the acute scans were kept.

The dataset (organized in BIDS format, bad quality / duplicate scans removed), was preprocessed with *fmriprep* (Esteban et al., 2019) on a supercomputer using *Apptainer* (formerly known as *Singularity*, Kurtzer et al., 2017).

Results included in this manuscript come from preprocessing performed using *fMRIPrep* 23.1.3 (Esteban et al. (2019); Esteban et al. (2018); RRID:SCR_016216), which is based on *Nipype* 1.8.6 (K. Gorgolewski et al. (2011); K. J. Gorgolewski et al. (2018); RRID:SCR_002502).

### Anatomical data preprocessing

The T1-weighted (T1w) images were corrected for intensity non-uniformity (INU) with `N4BiasFieldCorrection` (Tustison et al., 2010), distributed with ANTs (Avants et al., 2008, RRID:SCR_004757), and used as T1w-reference throughout the workflow. The T1w-reference was then skull-stripped with a *Nipype* implementation of the `antsBrainExtraction.sh` workflow (from ANTs), using OASIS30ANTs as target template. Brain tissue segmentation of cerebrospinal fluid (CSF), white-matter (WM) and gray-matter (GM) were performed on the brain-extracted T1w using `fast` (FSL, RRID:SCR_002823, Zhang et al., 2001). Volume-based spatial normalization to one standard space (MNI152NLin2009cAsym) was performed through nonlinear registration with `antsRegistration` (ANTs), using brain-extracted versions of both T1w reference and the T1w template. *ICBM 152 Nonlinear Asymmetrical template version 2009c* [Fonov et al. (2009), RRID:SCR_008796; TemplateFlow ID: MNI152NLin2009cAsym] template was selected for spatial normalization and accessed with *TemplateFlow* (23.0.0, Ciric et al., 2022).

### Functional data preprocessing

For each of the BOLD runs found per subject (across all runs), the following preprocessing was performed. First, a reference volume and its skull-stripped version were generated using a custom methodology of *fMRIPrep*. Head-motion parameters with respect to the BOLD reference (transformation matrices, and six corresponding rotation and translation parameters) are estimated before any spatiotemporal filtering using `mcflirt` (*FSL*, Jenkinson et al., 2002). BOLD runs were slice-time corrected 0.5 of slice acquisition range using `3dTshift` from *AFNI* (Cox & Hyde, 1997, RRID:SCR_005927). The BOLD time-series (including slice-timing correction when applied) were resampled onto their original, native space by applying the transforms to correct for head-motion. These resampled BOLD time-series will be referred to as *preprocessed BOLD in original space*, or just *preprocessed BOLD*. The BOLD reference was then co-registered to the T1w reference using `mri_coreg` (*FreeSurfer*) followed by `flirt` (*FSL*, Jenkinson & Smith, 2001) with the boundary-based registration (Greve & Fischl, 2009) cost-function. Co-registration was configured with six degrees of freedom. Several confounding time-series were calculated based on the *preprocessed BOLD*: framewise displacement (FD), DVARS and three region-wise global signals. FD was computed using two formulations following Power (absolute sum of relative motions, Power et al. (2014)) and Jenkinson (relative root mean square displacement between affines, Jenkinson et al. (2002)). FD and DVARS are calculated for each functional run, both using their implementations in *Nipype* (following the definitions by Power et al., 2014). The three global signals are extracted within the CSF, the WM, and the whole-brain masks. The head-motion estimates calculated in the correction step were also placed within the corresponding confounds file. The confound time series derived from head motion estimates and global signals were expanded with the inclusion of temporal derivatives and quadratic terms for each (Satterthwaite et al., 2013). The BOLD time-series were resampled into standard space, generating a *preprocessed BOLD run in MNI152NLin2009cAsym space*. First, a reference volume and its skull-stripped version were generated using a custom methodology of *fMRIPrep*. All resamplings can be performed with *a single interpolation step* by composing all the pertinent transformations (i.e. head-motion transform matrices, susceptibility distortion correction when available, and co-registrations to anatomical and output spaces). Gridded (volumetric) resamplings were performed using `antsApplyTransforms` (ANTs), configured with Lanczos interpolation to minimize the smoothing effects of other kernels (Lanczos, 1964). Non-gridded (surface) resamplings were performed using `mri_vol2surf` (FreeSurfer).

Many internal operations of *fMRIPrep* use *Nilearn* 0.10.1 (Abraham et al., 2014, RRID:SCR_001362), mostly within the functional processing workflow. For more details of the pipeline, see the section corresponding to workflows in fMRIPrep’s documentation.

### Copyright Waiver

The above boilerplate text was automatically generated by *fMRIPrep* with the express intention that users should copy and paste this text into their manuscripts *unchanged*. It is released under the CC0 license.

### Postprocessing of the functional data and generating connectivity matrices

Following a quality control of *fMRIPrep* outputs, 26 control subjects and 104 stroke subjects were qualified for further analysis. Each subject had 1-7 RS runs (112 subjects with 7 runs, 9 subjects with 6 runs, 4 subjects with 5 runs, 4 subjects with 3 runs, and 2 subjects with 2 runs), in total 875 runs for 130 subjects. Among 104 stroke subjects, 84 of them had manually drawn lesion masks available in the dataset. Regions that are most often damaged are the subcortical area, covering the thalamus and basal ganglia (Figure 1).

**Figure 1.**
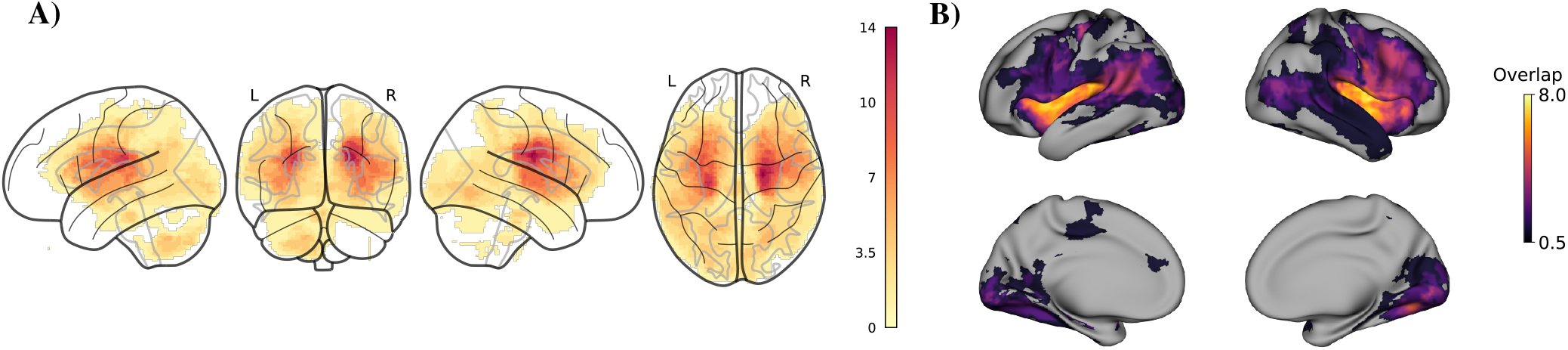
Profiles of ischemic stroke occurrence. **A)** Overlap of the 84 manually drawn lesion masks in the subcortical and cerebellar areas. Regions around the basal ganglia and thalamus show the most overlap, especially in the right hemisphere. The colorbar shows the number of subjects. **B)** Cortical projection of the lesion occurrences. Total number of subjects with cortical lesion occurrence is less than the total number of subjects with subcortical lesion occurrence.

Outputs of *fMRIPrep* were further cleaned from nuisance regressors using a custom script written in MATLAB. The 36P strategy without global signal (GS) regression described in Satterthwaite et al. (2013) was adopted to choose the regressors of no interest. Specifically, 6 motion regressors, mean signal from white matter and CSF, their first derivatives, power, and power of the first derivatives were chosen. In addition, mean frame-wise displacement and a linear trend were added to the model. These regressors were removed from the fMRI data using a multiple linear regression model and the residuals were saved. A band-pass filter between 0.01-0.1 was applied to the residuals.

Mean time series from GM were calculated using an improved version of the Schaefer-Yeo atlas that parcellates the cortex into 400 regions across 7 functional networks (Glen et al., 2021; Schaefer et al., 2018). The averaged time series were then correlated with each other, resulting in 400 x 400 correlation matrices for each run of each subject. Average correlation matrices across runs were calculated and normalized via Fisher’s Z transformation. This step generated one correlation matrix for each subject.

### Calculation of the temporal lag

The time lag of the BOLD signal was calculated in relation to the GS, which was derived from the GM mask (whole-brain GM mask for controls, and only the intact hemisphere for the stroke subjects). For each voxel on the cortex, the correlation between the BOLD signal and the time-shifted version of the GS signal between ±3 TR was calculated. The time-shift value that corresponds to the maximum absolute correlation was accepted as the time lag. The BOLD signal was then shifted to that many TRs to correct for the time lag.

400 × 400 correlation matrices were generated from the corrected time series using the same procedure for the non-corrected time series. We used the functional connectivity aberrancy to measure the effect of this correction on the connectivity matrices (Siegel, Snyder, et al., 2016). This method calculates the Z score of each node in the individual correlation matrices in reference to the mean connectivity value of the control subjects. For each subject (both control and stroke groups), mean aberrancy across the whole matrix was calculated. These mean values of groups were compared via a two-sample t-test before and after lag correction.

### Calculating the functional connectivity gradients and the reference topography

Functional connectivity gradients based on the correlation matrices were generated as described in Margulies et al. (2016) using the available functions in the *BrainSpace* MATLAB toolbox (Vos de Wael et al., 2020). In short, correlation matrices were thresholded to keep the top 10% of the connections. The normalized angle similarity of the sparse matrices was calculated. A non-linear dimension reduction technique, diffusion mapping, was applied to the similarity matrices with an alpha value of 0.5 for the manifold. This step generated 10 gradient values that explain the functional connectivity profile of each ROI and for each subject. Lastly, Procrustes alignment was used on the gradients to make them comparable. Since diffusion mapping can result in values in the opposite directions, a reference gradient computed based on the correlation matrix that came with *BrainSpace* was used. This procedure generated “gradient scores” that show the position of each region for 10 gradients. Only the scores of the first three gradients were kept for the following analyses. Gradients of the lag-corrected matrices were also calculated following the same procedure for the non-corrected data. Gradient scores of each ROI before and after lag correction were compared before and after lag correction within groups via within-samples t-test. After the evaluation of the aberrancy and gradient scores, lag-corrected data was used for further analyses. However, in order to quantify the effect of this correction, all analyses were repeated using the non-corrected data as well.

### Euclidean distance to the reference topography

The first 3 gradients (**G1, G2, G3**) constitute the orthogonal axes of a Euclidean space (i.e., 3D space) where the functional topography can be embedded. One of the main advantages of this approach is the considerably lower number of dimensions present in the data. Since this space is normed, we defined the position in this latent space as **x**_*s*,*r*_ = (*G*1_*s*,*r*_, *G*2_*s*,*r*_, *G*3_*s*,*r*_) for each subject *s* and ROI *r*. Then, for each ROI, a reference 3D organization was created by averaging the distribution over all control subjects 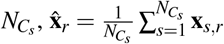 (which will be referred as reference topography). The Euclidean distance (ED) to the reference topography was calculated for every control and stroke subject as the *L*2 norm of the difference between the ROI’s and the reference positions, 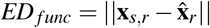. The ED to the 3D reference topography will be referred to as *ED*_*func*_, and was calculated using MATLAB’s *norm* function.

The mean *ED*_*func*_ across all subjects was calculated. This step generated 400 *ED*_*func*_ values for control and stroke groups separately. The mean *ED*_*func*_ values were compared via a two-sample t-test in order to check if both groups have equal mean distance to reference topography. The significance of the region-wise distances was calculated via a permutation procedure. The mean *ED*_*func*_ of the stroke group was subtracted from that of the control group, leaving a vector of 400 differences (one difference value for each ROI). The same difference vector was calculated 10000 times after randomly shuffling the group labels, calculating the mean *ED*_*func*_ for each group. This procedure created a null distribution of difference values for each ROI. The value of the real difference of an ROI was considered significant if its absolute value exceeded the 95th percentile value of its null distribution. To detect the source of the difference in *ED*_*func*_, 1 dimensional (1D) *ED*_*func*_ for every one of the three gradients was also calculated separately and compared across groups following the same procedure.

### Quantification of intrahemispheric functional connectivity profiles

In stroke patients, lesion lateralization is known to play an important role in behavioral and cognitive impairments. To study the functional connectivity profiles in intact and lesioned hemispheres independently we computed the connectivity gradients for intrahemispheric connections using the same methods. Importantly, out of 130 participants with lesions on the left side (*n* = 56) and on the right (*n* = 48) were treated separately, and intrahemispheric subsets of the (400 × 400) matrices were extracted for each patient. Intact and lesioned hemispheric functional connectivity gradients were then computed based on the similarity profiles of the (200 × 200) functional connections.

To examine how the functional reorganization differs in the right and left hemispheres in the case of lesion and no lesion, 3D and 1D *ED*_*func*_ for right and left hemispheres were compared within intact and damaged hemispheres. The mean *ED*_*func*_ across all subjects was compared between the right and left hemispheres separately. Additionally, a one-way ANOVA was used to compare the lateralization (right *ED*_*func*_ minus left *ED*_*func*_) of control, intact, and damaged hemispheres.

### Effect of lesion location on ED_func_

To examine whether the stroke lesion on close regions is related to similar functional reorganization profiles, the Jaccard similarity index (JI) between lesion masks of each subject and the correlation between their *ED*_*func*_ values were calculated. This step created 83 JI values and 83 correlation coefficients for each subject (there are 84 subjects with lesion masks). The values across subjects were pooled together, resulting in 3486 JI and correlation values. The correlation between the two sets was examined to investigate the similarity between lesion location and functional deviation patterns. The analysis was repeated with ED between centroids of the lesion masks (*ED*_*centroid*_) instead of JI, in case of 0 overlap between the masks.

### Relationship between ED_func_ and behavioral-structural measures

The behavioral examination includes five main batteries that consist of various tests (the full list of tests is available in Corbetta et al., 2015). These batteries measure the performance of the motor, linguistic, executive function, memory, and attentional capabilities. It has been shown that multiple tests could be clustered into a reduced set of factors to account for a large variance of functional abnormalities (Corbetta et al., 2015). Three big clusters were identified to account for 60% variance in the behavioral performances. The first big cluster accounted for 25% of the variance and consisted of language, verbal memory, and spatial memory performances. The second big cluster explained 23% of the variance and consisted of visual field bias, left body, and spatial memory factors. Lastly, the third big factor explained 17% of the variance and consisted of right body and attention shifting factors. For each one of the aforementioned domains, we considered the clustered factor as a sufficient predictor of the given behavioral deficit. *ED*_*func*_ was correlated with the big cluster scores to investigate the relationship between functional and behavioral abnormalities.

Anatomical ED to lesion (*ED*_*anat*_) was calculated for each subject whose lesion masks were available. After resampling the lesion masks to the resolution of the functional scans, coordinates of the voxels that are marked as “lesioned” were extracted using AFNI’s *3dmaskdump*. Then ED between a voxel in the parcellation map and every lesioned voxel was calculated. The minimum distance to the lesioned voxels was defined as the distance to the lesion. This procedure was repeated for every voxel in the parcellation mask. Correlation between *ED*_*anat*_ and *ED*_*func*_ were for each region was calculated.

The visualizations in this paper were generated using functions from *BrainSpace* (Vos de Wael et al., 2020), *BrainStat*

(Worsley et al., 2009), *SurfPlot* (Gale et al., 2021), and adaptation of code from Bayrak (2019).

## Results

### Demographics and quality control measures

No difference was found between the mean ages of the groups (number of control subjects = 26, mean age = 55.3 ± 12.94, number of stroke subjects = 104, mean age = 53.5 ± 10.73; *t*(128) = 0.73, *p* = 0.46) and sex distribution (stroke subjects = 54 male/50 female, control subjects = 16 male/10 female; 𝒳^2^ = 1.01, *p* = 0.3). Mean FD was higher in control subjects (*t*(128) = 4.81, *p* < 0.01). Variance and maximum values of FD did not show any difference between groups.

### Effect of lag correction on the connectivity matrices and gradients

Stroke subjects showed higher functional connectivity aberrancy compared to the control subjects before lag correction (*t*(127) = 2.43, *p* < 0.05). After the correction, there were no differences in terms of the functional connectivity strength between the groups (*t*(127) = 0.98, *p >* 0.05). Also, the mean functional connectivity of both groups was increased after the correction (Figure 2A-B).

**Figure 2.**
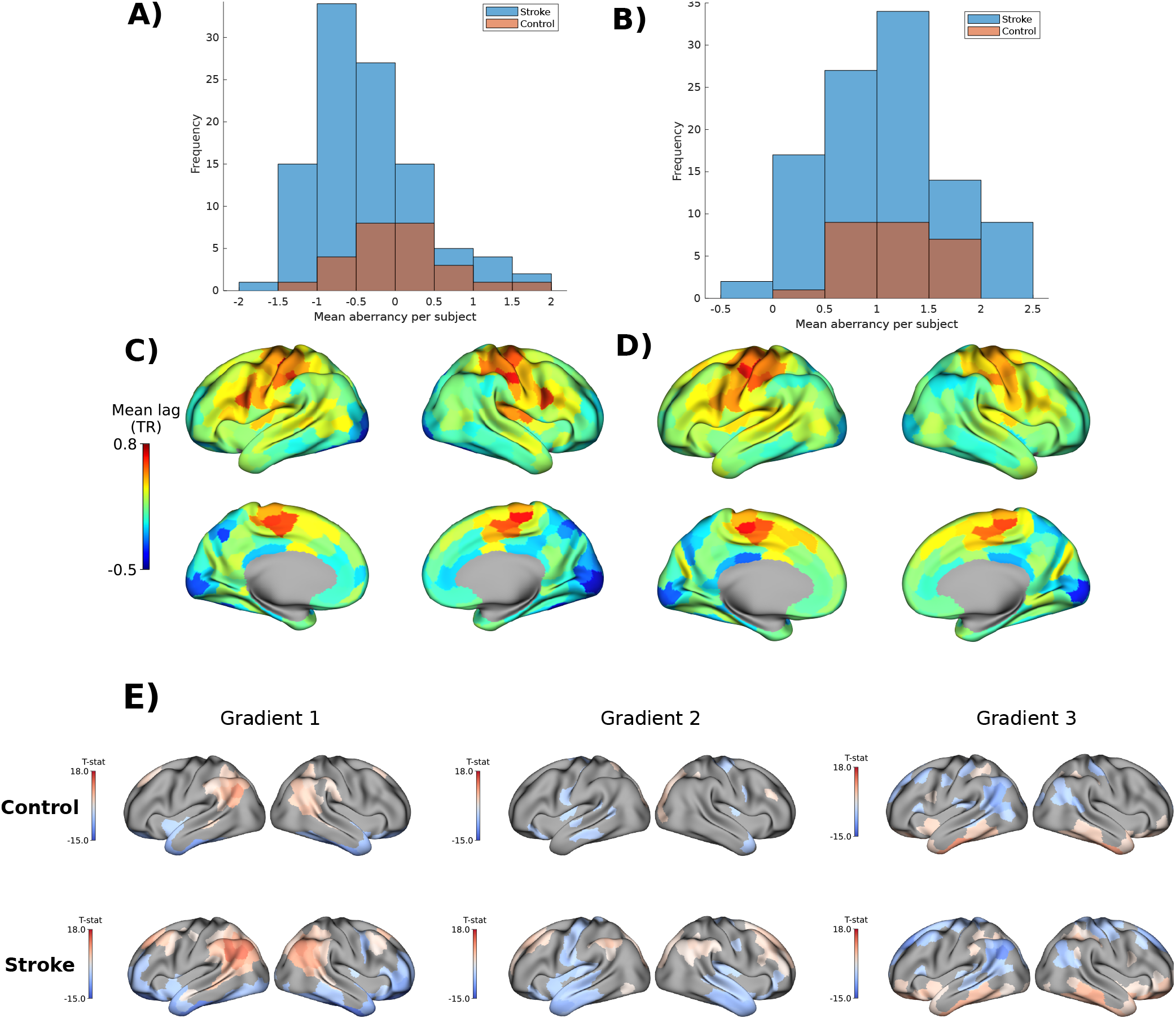
Effect of lag correction on connectivity matrices and gradient scores. **(A)** Before and, **(B)** after lag correction. Aberrancy is calculated in reference to the mean connectivity of control subjects before lag correction. It can be seen that stroke subjects show aberrancy before lag correction, but after lag correction, the data of the two groups align. The lag patterns of control and stroke groups can be seen in **(C)** and **(D)** respectively. **(E)**: Gradient scores of each region were compared before and after lag correction via a t-test, and p-values were corrected with FDR. Top row: Control group, Bottom row: Stroke group. Negative values depict higher gradient scores after correction. Stroke subjects show similar but larger differences in gradient scores after lag correction.

The mean lag maps in reference to GS showed a pattern on the inferior-superior axis (consistent with Erdoğan et al. (2016) and Tong et al. (2019); Figure 2C-D). The visual cortices have a negative delay, hence lagging behind the GS signal. On the contrary, somatomotor areas have a positive delay, meaning that they are followed by the GS. Mean, and variance of the lag values did not show any difference between the groups (*t*(798) = − 0.91, 1.17, *p* = 0.35, 0.24, respectively). However, maximum lag values were higher in stroke subjects, and minimum lag values were higher in controls (*t*(798) = − 13.72, 13.47, respectively, *p* < 0.001 for both, Supplementary Figure 2). In other words, stroke subjects showed more extreme lag values on both sides.

ROI-based comparison of the gradient scores before and after lag correction resulted in differences in both groups (*p* < 0.05, corrected with FDR, Figure 2E). In control subjects, higher gradient scores in temporal and frontal poles and lower scores on cuneus were observed for G1, higher scores in temporal poles and lower scores in visual cortices were observed for G2, and higher scores in cuneus and superior frontal gyri and lower score for temporal poles, frontal poles, and inferior temporal gyri for G3. For stroke subjects, these patterns were similar but more expanded, with the highest expansion occurring in the second gradient. ROI-based comparison was done to check the differences between control and stroke subjects, but the results no significant differences in gradient scores between two groups were found, neither before nor after lag correction.

Explained variances (i.e., scaled eigenvalues) also increased for G1 and G2 in stroke subjects after lag correction (*t*(103) = − 4.96, − 4.61, respectively, *p* < 0.05, corrected with FDR, Supplementary Figure 1). The mean explained variance increased by 0.97% ± 2 in G1 and 0.57% ± 1.24 in G2 (*p* < 0.05, corrected with FDR). When correlated with mean gradient scores, the mean lag pattern showed a very high correlation with G2 (*r* = 0.7), and lower with G1 (*r* = 0.25) and G3 (*r* = 0.2, *p* values < 0.001, FDR corrected).

To capture a similar nuance with *ED*_*func*_, for each subject, the difference between the individual lag value and the mean lag value from control subjects was calculated for each ROI. Then, the mean lag difference was correlated with mean *ED*_*func*_.

Noteworthy, control subjects showed almost zero correlation (*r* = 0.02, *p* = 0.6), whereas stroke subjects showed stronger correlation (*r* = 0.2, *p* < 0.001). The lag correction did not change those values (*r* = 0.02 and 0.21). Importantly, the results in the following sections are all based on the lag-corrected data.

### Connectivity gradients

Functional connectivity gradients revealed the functional similarity profile of the regions depending on how much variance they explain. On average, the first three gradients (G1, G2, G3) explained 41.63 % of the total variance for both groups (17.62%, 13.53%, and 10.48%, respectively). The following analyses were conducted on these gradients. There were no differences in the explained variance of the gradients between groups (explained variances for each gradient were compared via two-sample t-test, *p* > 0.05 for 10 gradients, corrected with false-discovery rate - FDR). The principal gradient G1 showed that brain activity is maximally organized between the unimodal/transmodal cortices. The second axis, or G2, showed the visual/somatomotor regions, and G3 showed the visual-default mode/dorsal attention-frontoparietal for both control and stroke subjects (Figure 3A). For each RS network, ROI-wise gradient scores in controls and patients were found to be statistically equivalent (*p* > 0.05, t-test, corrected with FDR). These results suggest that the functional topography remains largely unaltered in the early stages after the stroke occurs.

**Figure 3.**
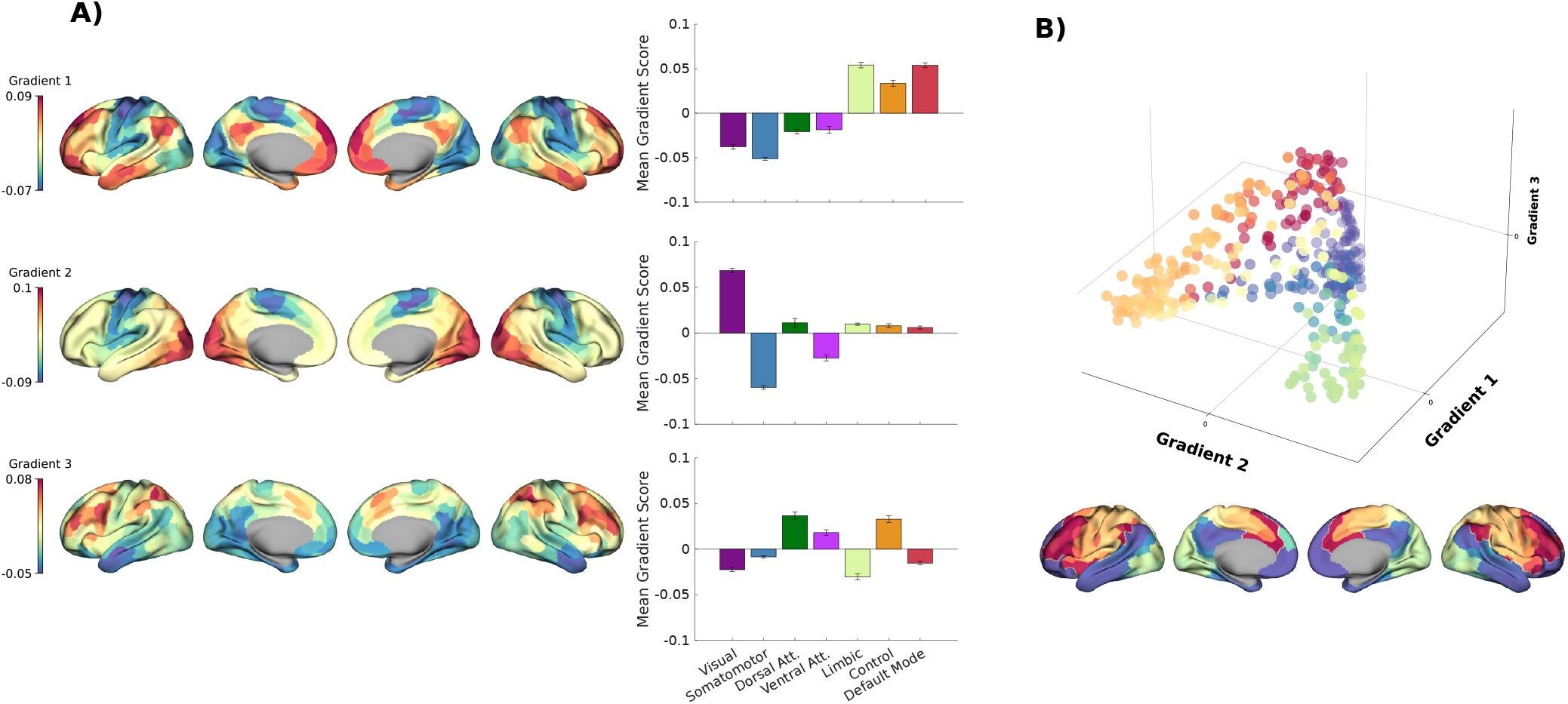
Reference topography from control subjects. **A)**: Projection of the mean gradient scores on the surface meshes (left) and their decomposition by 7 networks (right). Error bars depict 1 Standard Error of the Mean (SEM). **B)**: Reference topography in 3D space, and its projection on a surface mesh.

### Euclidean distance to functional organization

The mean 3D distribution to reference topography (*ED*_*func*_) was calculated for every ROI and every subject. The values of control and stroke subjects were compared via a two-tailed between-samples t-test. The results showed that stroke patients have significantly more mean *ED*_*func*_ compared to control subjects (*t*(798) = − 12.05, *p* < 0.001, Figure 4A). Permutation tests and the 7 networks decomposition revealed that the source of the difference is the somatomotor and ventral attention networks, more visible in the right hemisphere (Figure 4B-C). Comparison of 1D *ED*_*func*_ that were derived from each gradient separately showed that all 3 gradients contribute to this difference (T-stats of the G1, G2, G3: -4.01, -10.21, -6.31, respectively, p<0.01, Supplementary Figure 3). All 3 gradients showed high mean differences in the somatomotor network. In addition, G2 showed a high difference in the visual network and G3 showed a difference in the ventral attention and limbic networks. Taken together, stroke subjects showed larger functional deviation to the reference topography, specifically in somatomotor, ventral attention, and visual networks.

**Figure 4.**
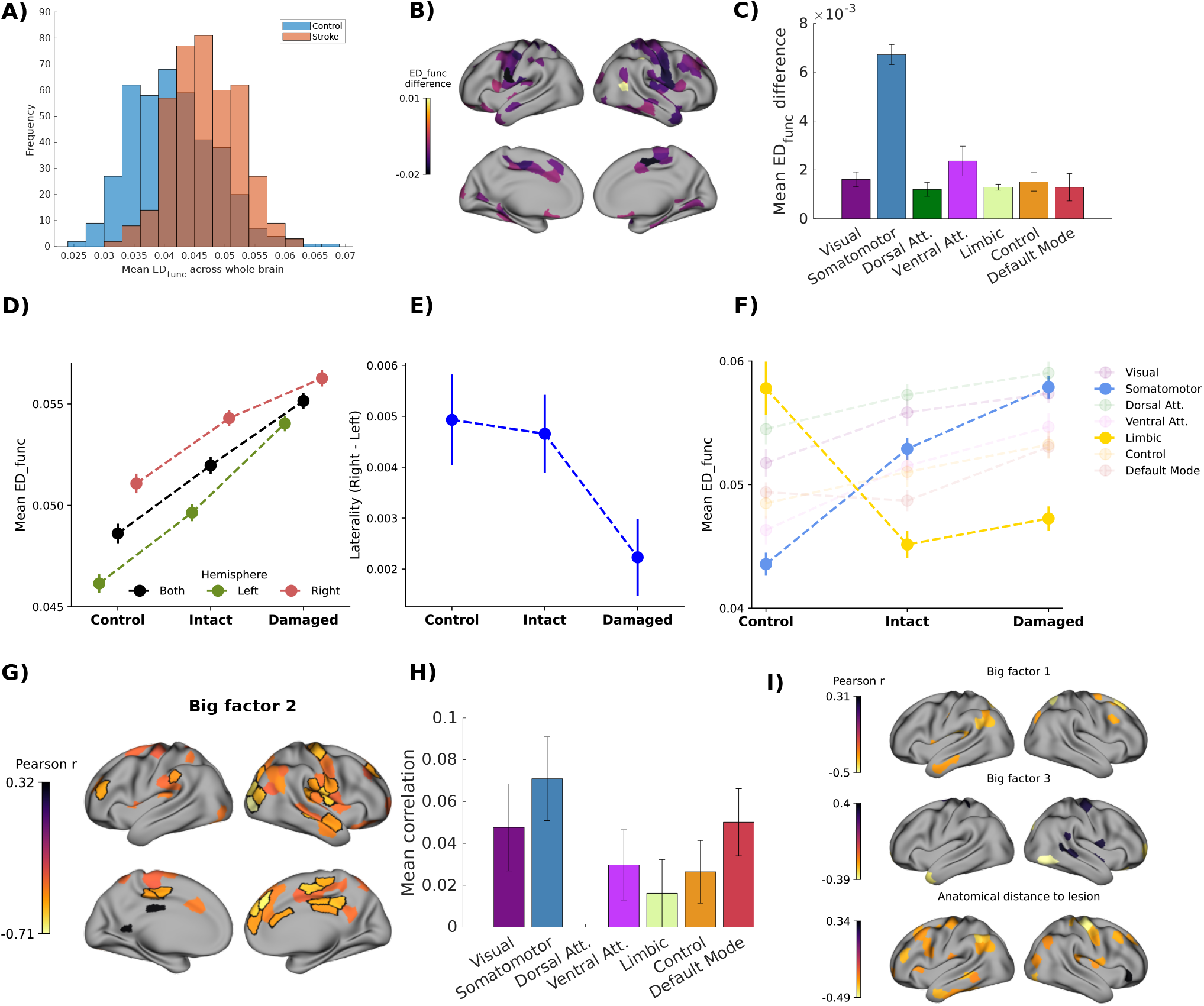
*ED* _*func*_ in control and stroke subjects, and its behavioral and anatomical correlates. **(A)** Histogram of *ED*_*func*_ between control and stroke subjects. **(B)** Significant regions-wise 3D *ED*_*func*_ differences (*p* < 0.05, corrected via permutation procedure). All differences favor stroke subjects, meaning that *ED*_*func*_ is larger in the stroke subjects for given regions. **(C)** Differences in *ED*_*func*_ across networks (error bars show mean *±* SEM). **(D)** *ED*_*func*_ difference across hemispheres of control subjects, and intact and damaged hemispheres of the stroke subjects. *ED*_*func*_ shows an increasing trend from control to damaged hemispheres (*F* = 3.39, *p* < 0.05). **(E)** Lateralization difference (calculated as *ED*_*func*_ of right hemisphere - left hemisphere) across control. intact, and damaged hemispheres. The difference between hemispheres is decreased in damaged hemispheres. **(F)** *ED*_*func*_ of the hemispheres across 7 networks. Functional deviancy in the somatomotor network increases in damaged hemispheres, while it decreases in the limbic network. **(G)**Regions that show correlation between *ED*_*func*_ and big factor 2 that is related to the visual field bias and left motor performance. (*p* < 0.05). Black contour indicates regions that survived multiple comparison correction through non-parametric permutation procedures. **(H)** shows the network composition of the regions that survived the multiple comparison correction. **(I)** Correlation between *ED*_*func*_ and other behavioral factors and *ED*_*anat*_ (*p* < 0.05, not corrected for multiple comparisons).

### Intrahemispheric connectivity profiles and lateralization in ED_func_

Both damaged and intact hemispheres showed higher *ED*_*func*_ in right hemisphere (damaged hemispheres: *t*(398) = − 2.81, *p* < 0.005, intact hemispheres: *t*(398) = − 5.72, *p* < 0.001). The difference is stronger in the intact hemispheres. For reference, the right and left hemispheres of the control subjects were also compared (*t*(398) = − 0.70, *p* = − 5.27). Lateralization differences among the control, intact, and damaged hemispheres were tested using a one-way ANOVA, revealing a significant model (*F* = 3.38, *p* = 0.035); Tukey’s post-hoc test indicated that lateralization differences were smaller in damaged hemispheres compared to control hemispheres (Figure 4D-E).

Repeating the same analysis using the 1D *ED*_*func*_ of each three gradient separately showed that the difference is specific to the second gradient in all three groups. In the control hemispheres, the difference for the 1st, 2nd, and 3rd gradients were respectively: *t*(398) = − 0.77, − 3.57, − 1.69, *p* = 0.43, 0.0004, 0.09; in the intact hemispheres: *t*(398) = − 1.83, − 3.42, − 2.68, *p* = 0.06, 0.0007, 0.007; and, in the damaged hemispheres *t*(398) = 0.10, − 2.92, − 0.08, *p* = 0.91, 0.003, 0.93. When the *ED*_*func*_ of the three groups were compared to each other across the whole brain, without separating the right and left hemispheres, an increasing trend was observed from control to damaged hemispheres (tested with one-way ANOVA, *F* = 46.15, *p* < 0.0005, corrected with Tukey’s post-hoc test, Figure 4D). Decomposition of the *ED*_*func*_ across 3 types of hemispheres showed that *ED*_*func*_ in somatomotor network and limbic network are the ones that change the most in damaged hemispheres, compared to the control ones (Figure 4F).

### Effect of lesion location on ED_func_

The correlation between JI, *ED*_*centroid*_, and *ED*_*func*_ showed *ED*_*func*_ and *ED*_*centroid*_ are weakly correlated (*r* = − 0.04, *p* < 0.05). The correlation becomes stronger when the subjects with right hemisphere damage are considered only (*r* = − 0.12, *p* < 0.005), and disappears for the subjects with left hemisphere damage (*r* = − 0.03, *p >* 0.05). The center of the lesion is weakly related to similar functional deviation patterns when the damage is on the right hemisphere (Table 1).

**Table 1.**
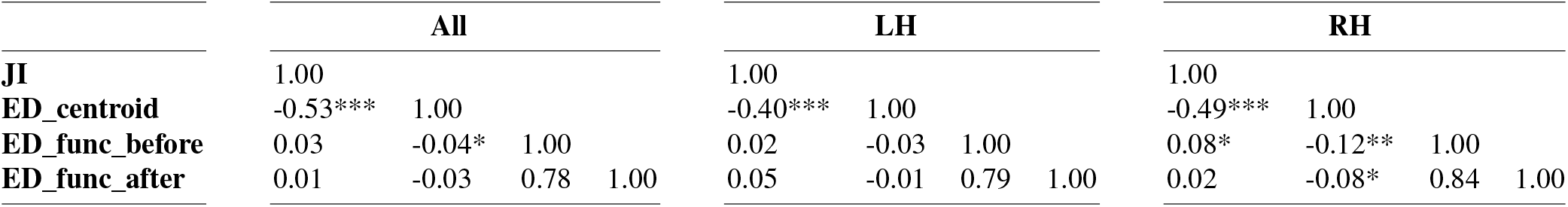
Correlation between Jaccard Index (JI), ED between centroids of the lesions (*ED*_*centroid*_), and *ED*_*func*_ before and after lag correction. across all subjects (All), subjects with right hemisphere damage (RH), and subjects with left hemisphere damage (LH) (* : *p* < 0.05, ** : *p* < 0.005, *** : *p* < 0.0005)

### Relationship between ED and behavioral-structural measures

Among the 3 big clusters of behavioral scores, only the second big cluster, which is driven by visual field bias and left body performance showed significant correlations with *ED*_*func*_ (*n* = 42, *p* < 0.05, corrected via FDR. Mean correlation across 32 significant regions: -0.51 *±* 0.06, Figure 4G-H). Regions that show the highest correlation were from somatomotor, visual, and default mode networks, mainly on the right hemisphere. Although the other two factors showed consistent patterns, their results did not survive FDR correction. Similarly, the correlation between *ED*_*func*_ and *ED*_*anat*_ showed a negative relationship in somatomotor and control networks (Figure 4I). However, the results did not survive FDR correction as well.

### Effect of lag correction on the results

The comparisons related to *ED*_*func*_ were repeated on the non-corrected data (Table 2). The explained variance of G1 and G2, the 1D *ED*_*func*_ of G2, and the lateralization of control, intact and damaged hemispheres increased after applying the lag correction. On the contrary, correlation between *ED*_*func*_ and mean lag difference as well as behavioral performance did not change. Lastly, the similarity between lesion location and functional reorganization decreased.

**Table 2.** Effect of lag correction on the results. Lag correction enhanced the *ED*_*func*_-related metrics, but did not change the behavioral and structural correlates. (* : *p* < 0.05). n/a stands for “not appplicable”.

| Measurement | Before correction | After correction | Difference |
| --- | --- | --- | --- |
| Explained variance of G1 and G2 of stroke subjects | 16%, 13% | 17.62%, 13.53% | t(103)= -4.96, -4.61 * |
| 1D $ED_{func}$ of G2 | t(798) = -7.9 | t(798) = -10.21 | t(399) = 4.63 * |
| $ED_{func}$ of R-L intact hemispheres | t(398) = -3.43 | t(398) = -5.72 | t(398) = 3.43 * |
| $ED_{func}$ of R-L damaged hemispheres | t(398) = -0.06 | t(398) = -2.81 | t(398) = 4.47 * |
| $ED_{func}$ of R-L control hemispheres | t(398) = -0.7 | t(398) = -5.51 | t(398) = 6.75 * |
| 3D $ED_{func}$ between stroke and control subjects | t(798) = -10.1 | t(798) = -12.05 | t(399) = 1.86, p = 0.06 |
| Correlation between mean lag diff. And $ED_{func}$ | r = 0.02, 0.2 | r = 0.02, 0.21 | n/a |
| Behavioral correlates of $ED_{func}$ | n/a | n/a | n/a |
| Correlation between $ED_{func}$ and $ED_{centroid}$ | r = -0.12 | r = -0.08 | n/a |

## Discussion

The first aim of this study was to identify the effect of temporal lag correction on the connectivity gradients. It has been reported that lag correction improves the connectivity strength for various metrics (Erdoğan et al., 2016; Siegel, Snyder, et al., 2016; Tong et al., 2019). However, to our knowledge, no study up to date investigated the influence of the lag correction on the functional connectivity gradients. Lag correction changed the gradient values for the control subjects in all 3 gradients. Although the regions that show different scores vary across gradients, in all three of them, a change on the superior-inferior axis is observed. From this aspect, it can be said that time lag correction “fixes” the data on the temporal domain, decreasing the confounding effect of the blood arrival time. A similar phenomenon was observed in Erdoğan et al. (2016), where the authors reported an increased explained variance in functional connectivity after time lag correction, along with better-aligned residuals with respect to the global signal. Considering the cerebral blood circulation, which goes from inferior to superior regions, it could be expected to observe differences in this axis. A similar pattern with a larger extent was revealed for stroke subjects, which means they benefited more from the correction. The limbic network, which is located on the first end of this inferior-superior axis, showed the most pronounced difference after lag correction for stroke subjects (Figure 2). The largest difference in gradient scores between control and stroke subjects was observed in the second gradient, potentially due to the high spatial correlation between the blood arrival map and the second gradient map. This suggests that the functional organization in the second gradient may be related to blood arrival time.

The second aim of this study was to identify the effect of acute ischemic stroke on the functional organization of the brain. The connectivity gradients were used as a metric because of their ability to decompose the functional connectivity matrices into 3D distribution, which makes it possible to track functional changes in each region of each subject, regardless of the location and the extent of the stroke lesion. To measure functional reorganization we computed the Euclidean distance between the gradient profiles and a reference profile derived from the control population. Within this framework, lag correction amplified these distances, both between groups and between hemispheres. Differences between control and stroke subjects before correction became larger; that is, functional deficiency in the second gradient and the right hemisphere increased.

Although the gradient scores did not show any differences at the group level, a comparison of the functional deviation values showed that stroke is related to large functional deviations, primarily in the somatomotor network, followed by the visual and ventral attention networks. The changes related to these networks have been reported before and found related to behavioral dysfunctions such as motor problems, aphasia, and attentional deficits (Carter et al., 2010; Corbetta & Shulman, 2002; He et al., 2007; Siegel, Ramsey, et al., 2016). The functional deviation in our study was correlated to left motor and visual attention performance as well. Considering the consistency with previous reports, it can be said that functional deviation, as defined with respect to a reference landscape in gradient space, is a valid estimator of the impact of the stroke on brain function. Lastly, regions with significantly altered gradient profiles that correlated with behavioral scores also exhibited a relationship with anatomical distance to the lesion. The further away they are from the lesion, the less functional deviation they exhibit. The only study that investigated the effect of ischemic stroke on connectivity gradients up-to-date (Bayrak et al., 2019) also points out the fact that changes in the connectivity gradients are related to the location of the lesion. Taking all these associations into account, only a set of regions, mainly in the right somatomotor area, are sensitive to the stroke lesion.

One possible explanation for the concentration of the sensitive regions on the right hemisphere and somatomotor network could be the slightly higher number of overlapped lesions on the right hemisphere, but the distribution of the lesions was fairly symmetric when considering absolute numbers. Instead, the distribution of the lesions on the gray matter closely resembled the pattern of the functional deviations (Figure 1). It could be said that functional deviation occurs mostly on the hemisphere that is ipsilateral to the lesion. However, if this were true that would mean that functional deviation would solely localized in the regions closely surrounding the aformentioned cortical projections. When we isolated subjects according to lesion side (i.e., left/right hemisphere), functional reorganization was not side-especific (Figure 5). For subjects suffering from a left hemisphere lesion, displayed more homotopic profiles of functional deviation, instead, right-side strokes increased functional deviations mostly on the ipsilateral hemisphere. This distinction shows that the accumulation of sensitive regions on the right side cannot be only related to the distribution of the lesions in the present dataset.

**Figure 5.**
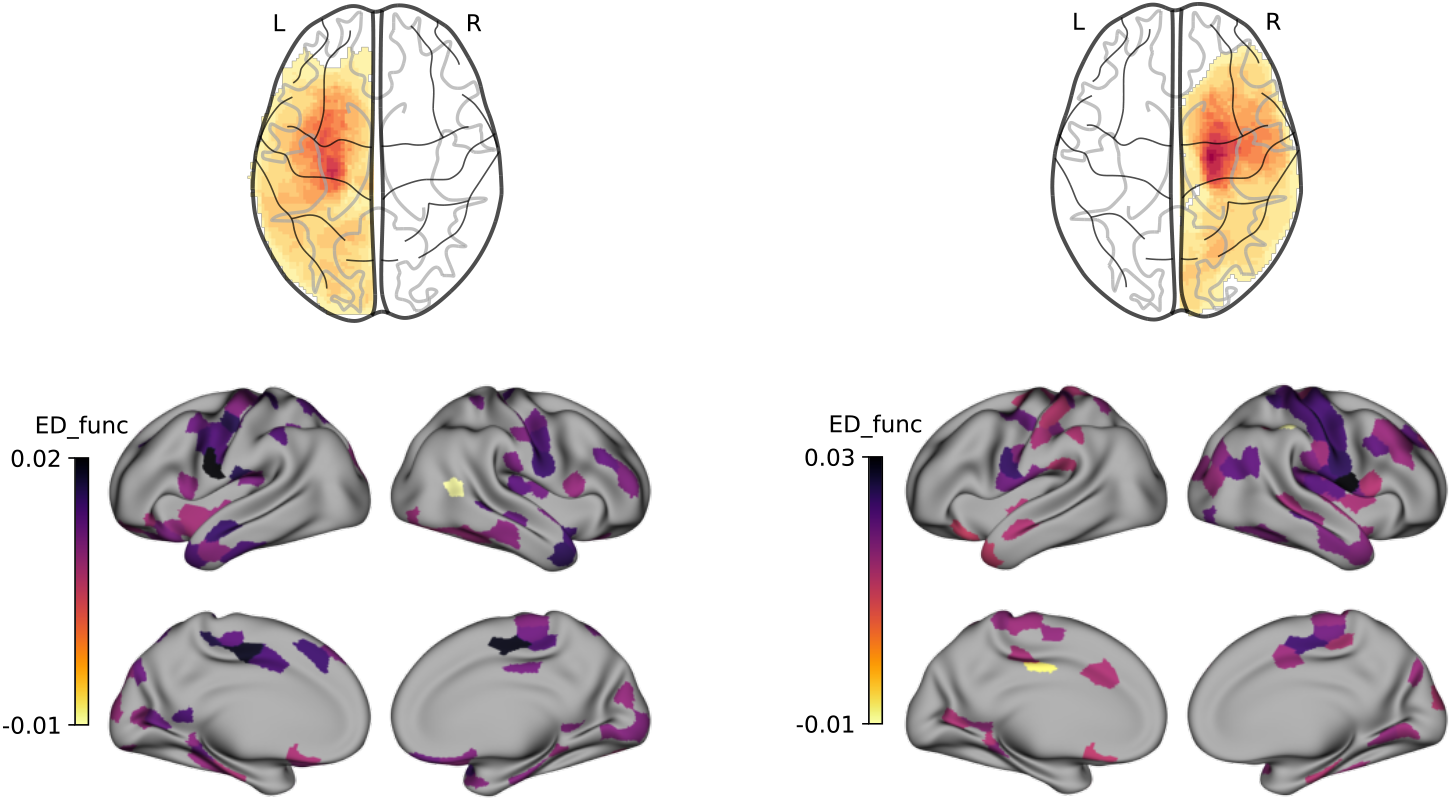
Effect of the lesioned hemisphere on the functional topography. Whole-brain functional connectivity gradients show different patterns of *ED*_*func*_ in stroke patients with left (*n* = 56) versus right hemisphere (*n* = 48) lesions. Left hemisphere lesions result in more distributed reorganization patterns, while right hemisphere lesions produce more localized effects. ROI-based *ED*_*func*_ analysis is presented after FDR correction.

An alternative explanation could be that the functional integrity of the right hemisphere is more susceptible to injuries due to language lateralization (Hodgson et al., 2014), and the highlighted sensitive regions are driving this hemispheric difference and behavioral deficits. This claim can be supported by the lateralization findings that showed similar deviation profiles when the damage occurs on the right hemisphere and the left hemisphere is preserved. In other words, functional reorganization is more consistent and similar when the left hemisphere is intact.

Lateralization may be understood in several ways, the most typical one being a patent asymmetry between the contribution of homotopic regions to a given brain function (Xu et al., 2024). A stroke is usually located in one hemisphere, hence, we studied whether functional gradients computed within hemispheres reacted in a similar way or rather acted independently with respect to differences present in the control population (New et al., 2015; Tang et al., 2016; Urbin et al., 2014; Yourganov et al., 2021). Interestingly, in control subjects, the right hemisphere was located further apart from the reference topography, indicating natural hemispheric asymmetry. Moreover, this remained unchanged for the intact hemispheres of the patients. However, when considering the lesioned hemisphere, this lateralization significantly decreased, suggesting that a stroke on the left hemisphere damages the functional organization to a larger extent. From another perspective, although the overall functional deviancy increases in the case of stroke, the lateralization profile is preserved if the left hemisphere is spared. Lag correction made these hemispheric differences more comparable between control and intact sides (Table 2).

The main source of the functional difference was the second gradient, where the brain function is organized on the somatomotor-visual axis. This major change in the axis that is aligned between the unimodal functions is consistent with the fact that stroke subjects mainly suffer from a decrease in the performance of the unimodal functions, whereas the higher-order functions are less affected by the damage.

Whether the deviation compensates for stroke damage or results from it remains uncertain. The extent of deviation may reflect the degree of functional plasticity compensating for anatomical damage, or conversely, the level of dysfunction. The direction of the correlation between behavioral performance and deviation can help clarify this. We found that correlations between functional deviation and behavioral scores were almost always negative, indicating that better behavioral performance (more similar to that of healthy controls) corresponds with smaller deviations. This suggests that the deviation reflects the degree of functional deficit, and this deficit is enhanced after lag correction procedures.

Extreme lag values of stroke subjects and the increased explained variance after the correction also confirm that the time lag indicates the hemodynamic deficits of the stroke subjects. One difference that disappeared after lag correction is the similarity between functional deviation and the proximity between lesion locations. Lag correction decreased the already-low correlation between these two factors. Hence, it can be said that correcting for hemodynamic lags is useful to clean data from structure-related noise for stroke subjects.

It could be expected to have a smaller difference between groups after correction since the BOLD signal is fixed to the same time domain. On the contrary, the functional deviancy difference is more visible. Therefore, the difference in the BOLD characteristics, at least in the time domain, can be considered noise. Another point of view could be that the new BOLD signal after the stroke may reflect the real neural activity, that happened due to plasticity and should be accepted as a whole while comparing the data between groups. For example, Siegel, Ramsey, et al. (2016) argue that although lag correction improves the connectivity, it is not suggested because there is no way to fix the other differences related to the frequency domain. Lag correction might complicate those confounding even more. However, our findings (also theirs, Siegel, Snyder, et al., 2016) show that this is a necessary step, and the issues in the frequency domain should be handled separately. Inspiration could be drawn from the literature studying brain tumors, where it has been found that local and global shifts of the spectrum carry prognostic (Park et al., 2023) and network (Falcó-Roget et al., 2024) information.

### Limitations and further considerations

The number of healthy controls in the dataset remains short compared to the number of stroke subjects. Using a surface-based approach limited the ability to calculate the geodesic distance to the lesion, which could be a better indicator than the anatomical Euclidean distance. Using disconnectome maps instead of anatomical distance can increase the overall anatomical information as it was shown that it is a better estimator of stroke outcome. Special attention can be paid to the somato-cognitive action network when examining the motor cortex. In addition to correcting BOLD in the time domain, correcting the frequency domain may lead to new diagnostics and improve the data quality in the future.

### Conclusions

Connectivity gradients are a valid method to examine the effect of ischemic stroke due to their low-dimensional representation of functional organization. This method highlights the sensitive regions that are placed mostly on the right hemisphere and on visual-sensorimotor axis. Time lag correction is a suggested denoising method for processing fMRI data, especially if the sample includes ischemic stroke subjects. Correction of the time lag highlighted the differences between groups, especially for the second gradient.

## Acknowledgments

Authors thank Prof. Maurizio Corbetta for sharing the dataset and Dr. Sophie Valk for their valuable feedback and suggestions. The publication was created within the project of the Minister of Science and Higher Education “Support for the activity of Centers of Excellence established in Poland under Horizon 2020” on the basis of the contract number MEiN/2023/DIR/3796, and has received funding from the European Union’s Horizon 2020 research and innovation programme under grant agreement No 857533. This publication is supported by Sano project carried out within the International Research Agendas programme of the Foundation for Polish Science, co-financed by the European Union under the European Regional Development Fund. This research was supported in part by the PLGrid infrastructure. Computations have been partially performed on the ARES supercomputer at ACC Cyfronet AGH.

## Author Contributions

**Cemal Koba:** Conceptualization; Data curation; Formal analysis; Investigation; Methodology; Validation; Visualization; Writing - original draft; and Writing - review & editing. **Joan Falcó-Roget:** Conceptualization; Data curation; Investigation; Methodology; Validation; Visualization; Writing - original draft; and Writing - review & editing. **Alessandro Crimi:** Conceptualization; Resources.

## Data and code availabilty

Data deposition: Neuroimaging and neuropsychological data at https://cnda.wustl.edu/app/template/Login.vm. The code used to transform the data into BIDS format can be found at https://github.com/JoanSano/BIDS_Constructor. The code used to compute and analyze the functional connectivity gradients can be found at https://github.com/KobaCemal/StrokeGradients.

## Supplementary Material: Effect of acute ischemic stroke on functional connectivity gradients

**Supplementary Figure 1.**
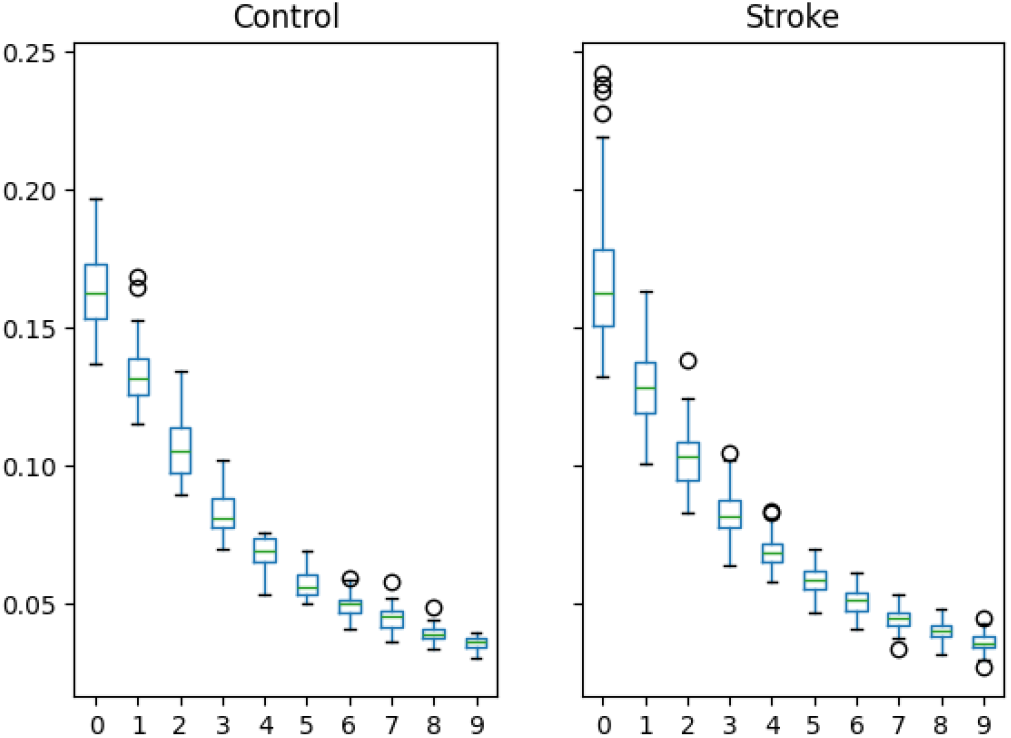
Scaled eigenvalues of the 10 gradients for control and stroke subjects.

**Supplementary Figure 2.**
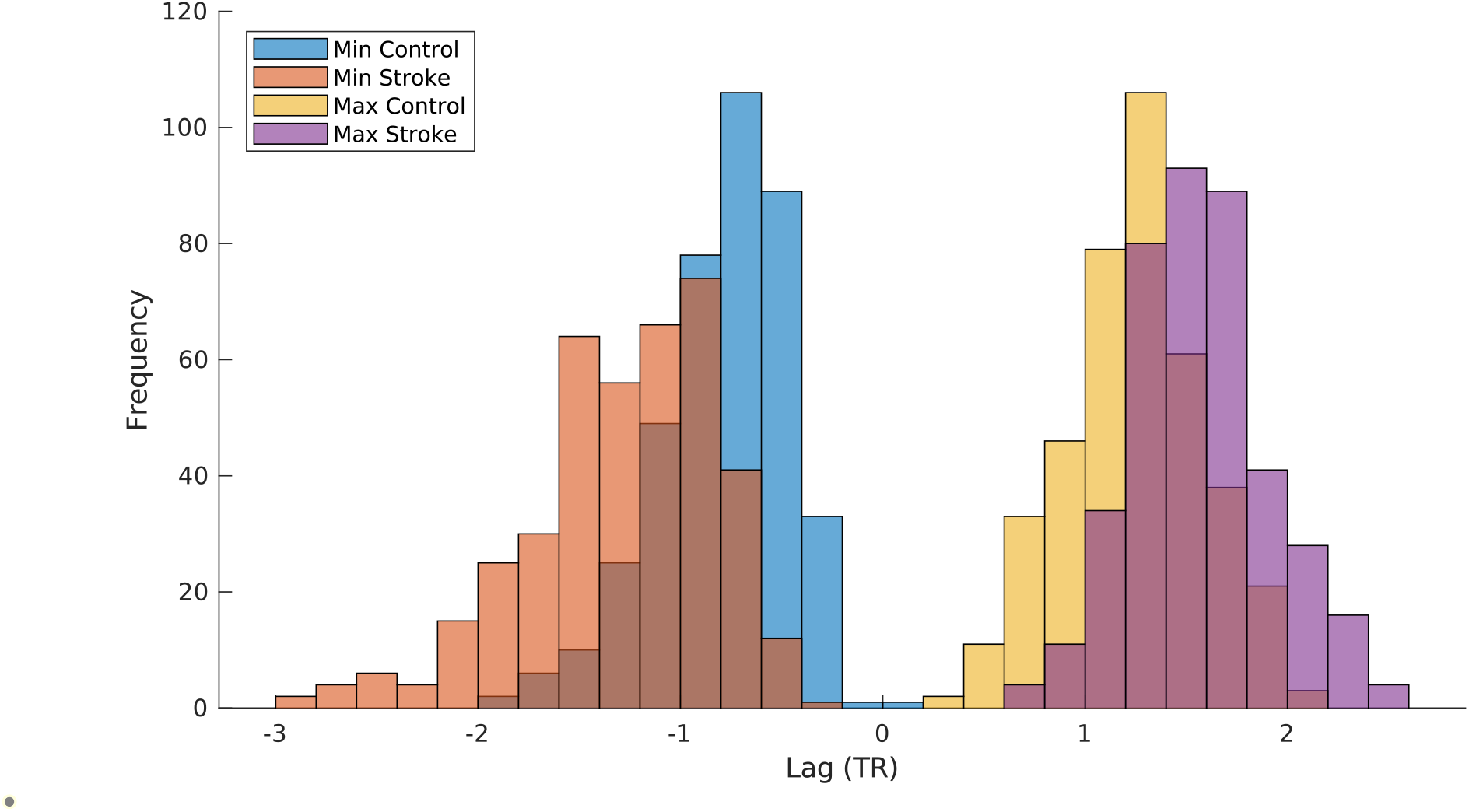
Minimum and maximum lag values of control and stroke subjects.

**Supplementary Figure 3.**
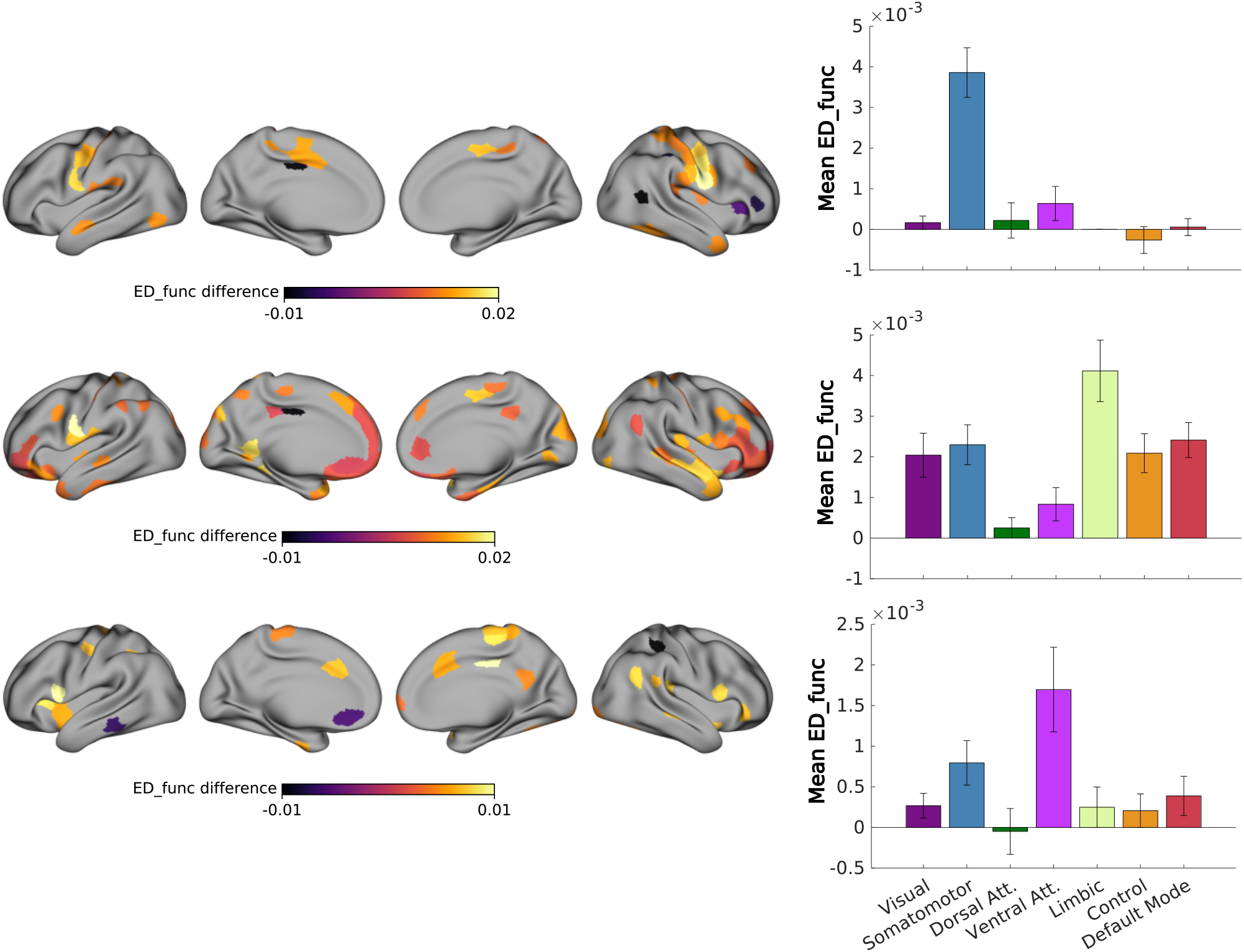
Significant regions-wise 1D *ED*_*func*_ differences (p < 0.05, corrected via permutation procedure).

